# Visualizing multiple inter-organelle contact sites using the organelle-targeted split-GFP system

**DOI:** 10.1101/290114

**Authors:** Yuriko Kakimoto, Shinya Tashiro, Rieko Kojima, Yuuki Morozumi, Toshiya Endo, Yasushi Tamura

**Author notes:** Correspondence and requests for materials should be addressed to Y.T.

## Abstract

Functional integrity of eukaryotic organelles relies on direct physical contacts between distinct organelles. However, the entity of organelle-tethering factors is not well understood due to lack of means to analyze inter-organelle interactions in living cells. Here we show that the split-GFP system can be used to observe organelle contact sites *in vivo*. We observed punctate GFP signals from the split-GFP fragments targeted to any pairs of organelles among the ER, mitochondria, peroxisomes, vacuole and lipid droplets in yeast cells, which suggests that these organelles form contact sites with multiple organelles simultaneously. Importantly, split-GFP signals in the overlapped regions of the ER and mitochondria were mainly co-localized with ERMES, an authentic ER-mitochondria tethering structure, suggesting that split-GFP assembly depends on the preexisting inter-organelle contact sites. We also confirmed that the split-GFP system can be applied to detection of the ER-mitochondria contact sites in HeLa cells. We thus propose that the split-GFP system is a potential tool to observe and analyze inter-organelle contact sites in living yeast and mammalian cells.

---

One of the functional and structural hallmarks of eukaryotic cells is the presence of complex membrane structures called organelles. Organelles enclose specific sets of enzymes and maintain spatial individuality to fulfill each organelle’s specialized functions. However, these classic views on organelle functions and structures have been radically challenged by the findings of physical connections between distinct organelles. These inter-organelle connections or contacts could allow exchange of organelle constituents like proteins, lipids, and metabolites as well as information between organelles. For example, the ERMES (endoplasmic reticulum (ER)–mitochondria encounter structure) complex, which consists of four core components, Mmm1, Mdm10, Mdm12 and Mdm34 (Mmm2), constitutes the best-characterized tethering structure, which directly connects the ER membrane and mitochondrial outer membrane (MOM) in yeast cells^1^. ERMES form clusters, which can be observed as a limited number of stable dots around the overlapping regions between mitochondria and the ER tubules under a fluorescent microscope by tagging the ERMES core components with a fluorescent protein such as GFP^2–6^. This indicates that specific regions of the mitochondrial surface form direct contact sites with the ER membrane through ERMES. *In vivo* pulse-chase experiments and *in vitro* reconstitution assays revealed that the ERMES complex facilitates phospholipid transport between these organelles at the ER-mitochondria contact sites, suggesting the functional importance of ERMES^1,7^. In addition to the ERMES complex, EMC (conserved ER membrane protein complex) present in the ER membrane was shown to interact with Tom5, which is a subunit of the mitochondrial outer membrane protein translocator, the TOM complex, and was suggested to be important for phospholipid transfer from the ER to mitochondria^8^.

vCLAMP (vacuole and mitochondria patch) was identified as a mitochondria-vacuole tethering structure, which appears functionally redundant with ERMES; (1) loss of vCLAMP component such as Vps39 increases the number of ERMES dots in a cell; (2) simultaneous loss of ERMES and vCLAMP components is synthetic lethal for yeast cells; (3) overexpression of Vps39 rescues the defective cell growth due to the loss of an ERMES component; (4) loss of ERMES extensively expands the vCLAMP region under microscope^9,10^. The functional redundancy between ERMES and mitochondria-vacuole contact sites was also supported by the findings that expression of a dominant mutant of an endosomal protein Vps13, which is localized to vacuole-mitochondria and vacuole-nucleus contact sites, can compensate the absence of ERMES^11^. Recently, Vps13 was shown to interact with a multi-spanning MOM protein Mcp1, the gene of which was identified as a multi-copy suppressor for ERMES components, and to contribute to recruitment of the vacuole membrane close to the MOM^12–14^. In addition to the ER–mitochondria and vacuole–mitochondria contacts, peroxisomes were reported to form contact sites with mitochondria and the ER through Pex11-ERMES interactions^15,16^.

A number of proteins have been found to form nuclear-vacuole contact sites called the NVJ (nuclear-vacuole junction)^17–20^. First, a nuclear membrane protein Nvj1 was found to function as a bridge between the nucleus and vacuole through direct binding to a vacuolar protein Vac8^21^. Formation of the NVJ may be critical for selective degradation of a part of the nucleus under a starvation condition^22^. Similar to other organelle contact sites, the NVJ appears to be involved in the lipid homeostasis because several proteins that are known to participate in lipid metabolism are enriched at the NVJ^17,18^. For example, an oxysterol-binding protein (OSBP) Osh1 and a SMP-domain containing protein Nvj2, both of which could function as lipid transfer proteins, are enriched at the NVJ^23,24^. The enoyl-CoA reductase Tsc13 and a phosphatidic acid phosphatase Pah1 are also enriched at the NVJ and to play key roles in the sphingolipid and phospholipid metabolism, respectively^22,25^. Furthermore, a StART-like domain containing protein Ltc1/Lam6 is present not only at the NVJ but also at the contact sites among the ER, mitochondria and vacuole and could be responsible for the sterol-transport^26,27^.

Lipid droplets (LDs) are related to many aspects of lipid metabolism including storage of neutral lipids. LDs were reported to interact with several other organelles such as the ER, vacuole, mitochondria, and peroxisomes although organelle-tethering factors associated with LDs are largely unknown^28,29^.

These findings regarding inter-organelle contact sites collectively expanded our understanding of how different organelles are coordinated to respond to cellular demands, yet they raised a number of new questions. First, it is not clear at the moment whether all the organelle contact sites have already been identified or not. For example, we do not know if there is any contact site between the vacuole and peroxisomes. While inter-organelle contact sites are often redundant, it is not obvious how much each organelle-tethering structures contribute to the functions and formation of the entire organelle contact sites. Furthermore, it remains to be revealed how formation of different organelle contact sites are coordinated and regulated to achieve optimized logistics of lipids, metabolites and so on among different organelles.

As a first step toward addressing these issues, here we evaluated the split-GFP system as a tool to detect inter-organelle contact sites. The split GFP system is a combination of the GFP fragments containing β-strands 1-10 and β-strand 11 of GFP, which can spontaneously assemble with each other to form a complete β-barrel structure of GFP and emit GFP fluorescence^30,31^. We confirmed that split-GFP fragments, each of which was expressed on the ER and mitochondria surface separately in yeast cells, gave rise to GFP signals in a granular pattern under fluorescent microscope. These GFP signals were found to co-localize with ERMES, the ER-mitochondria contact sites. This indicates the potential ability of split-GFP as a tool to detect pre-existing organelle contact sites. Interestingly, the assembled split-GFP signals were observed mainly as discrete foci between all pairs of organelles among the ER, mitochondria, vacuole, peroxisomes, and LDs, suggesting that each organelle forms contact sites with limited areas of multiple different organelles simultaneously. Moreover, we confirmed that the split-GFP system could work for detecting the ER-mitochondria contact sites in human HeLa cells. Thus, the present study shows that the split-GFP system offers a potential tool to detect and analyze inter-organelle contact sites in yeast cells as well as mammalian cells, and hopefully to screen for unidentified tethering factors between organelles and/or their regulator proteins.

## Results

### Mitochondria-ER contact sites can be detected by the split-GFP system

We asked here whether the split-GFP system, which is generally used to detect protein-protein interactions^30,31^, can be applied to visualization of contact sites between multiple organelle pairs, like the split-Venus system, which was previously used to visualize cellular inter-membrane contact sites^8,23^. Although the split-Venus system was tested for detecting only the pairs between the ER and plasma membranes^23^ and between the ER and mitochondrial membranes^8^, we made a more thorough survey for detection of inter-organelle contact sites by using the split-GFP-fusion proteins summarized in Fig. 1. Even when separately expressed and targeted to different organelle membranes, the split-GFP fragments, the first ten β-strands (GFP1-10) and the last (11th) β-strand (GFP11), are expected to diffuse in each organelle membrane and to assemble with each other at membrane contact sites, resulting in emission of GFP fluorescent signals (Fig 2A).

**Figure 1:**
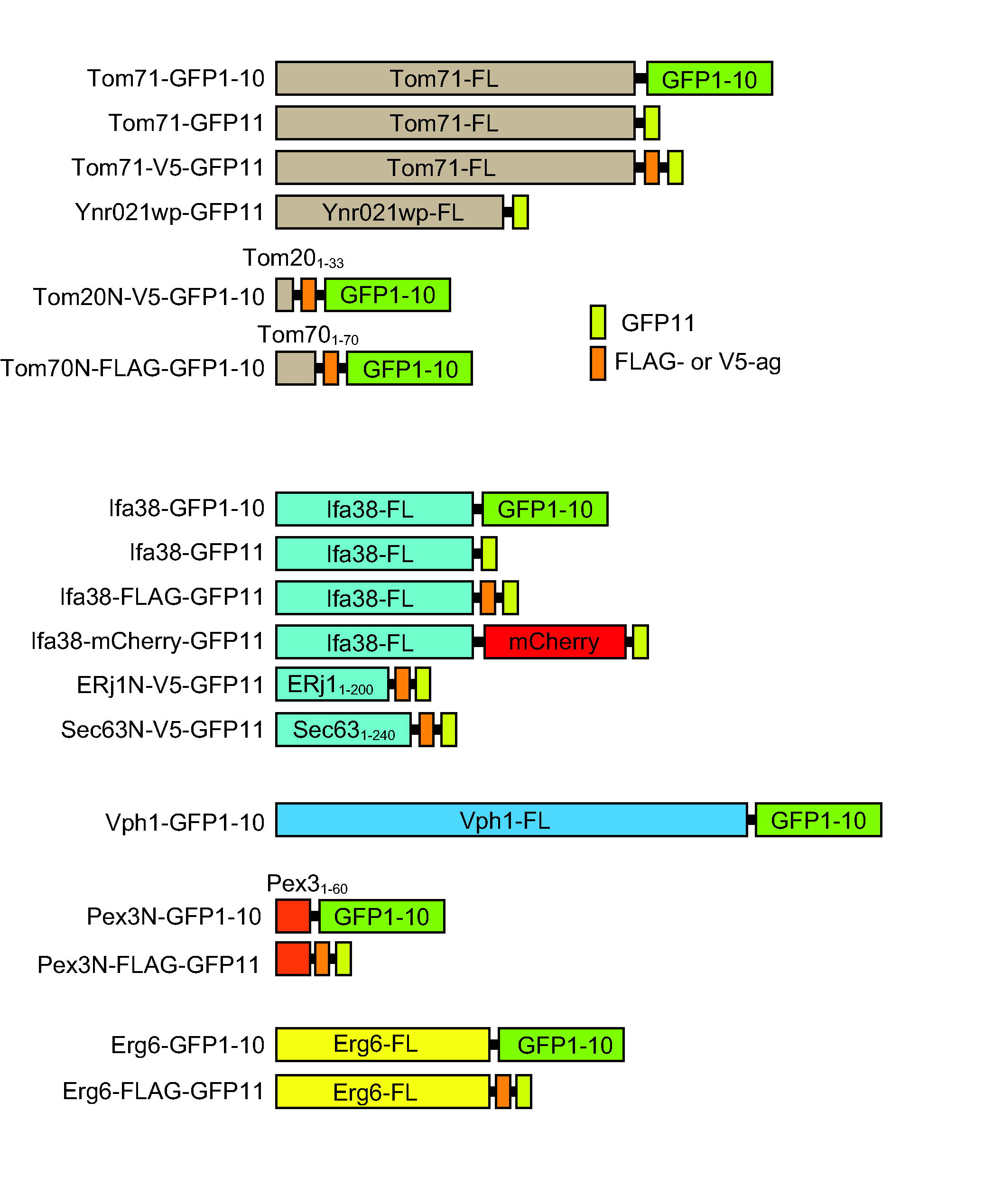
Schematic diagram of the split-GFP-fusion proteins used in this study. FL represents the full-length protein. Tom20_1-33_, Tom70_1-70_, ERj1_1-200_, Sec63_1-240_ and Pex3_1-60_ mean the N-terminal segments of the indicated proteins.

**Figure 2:**
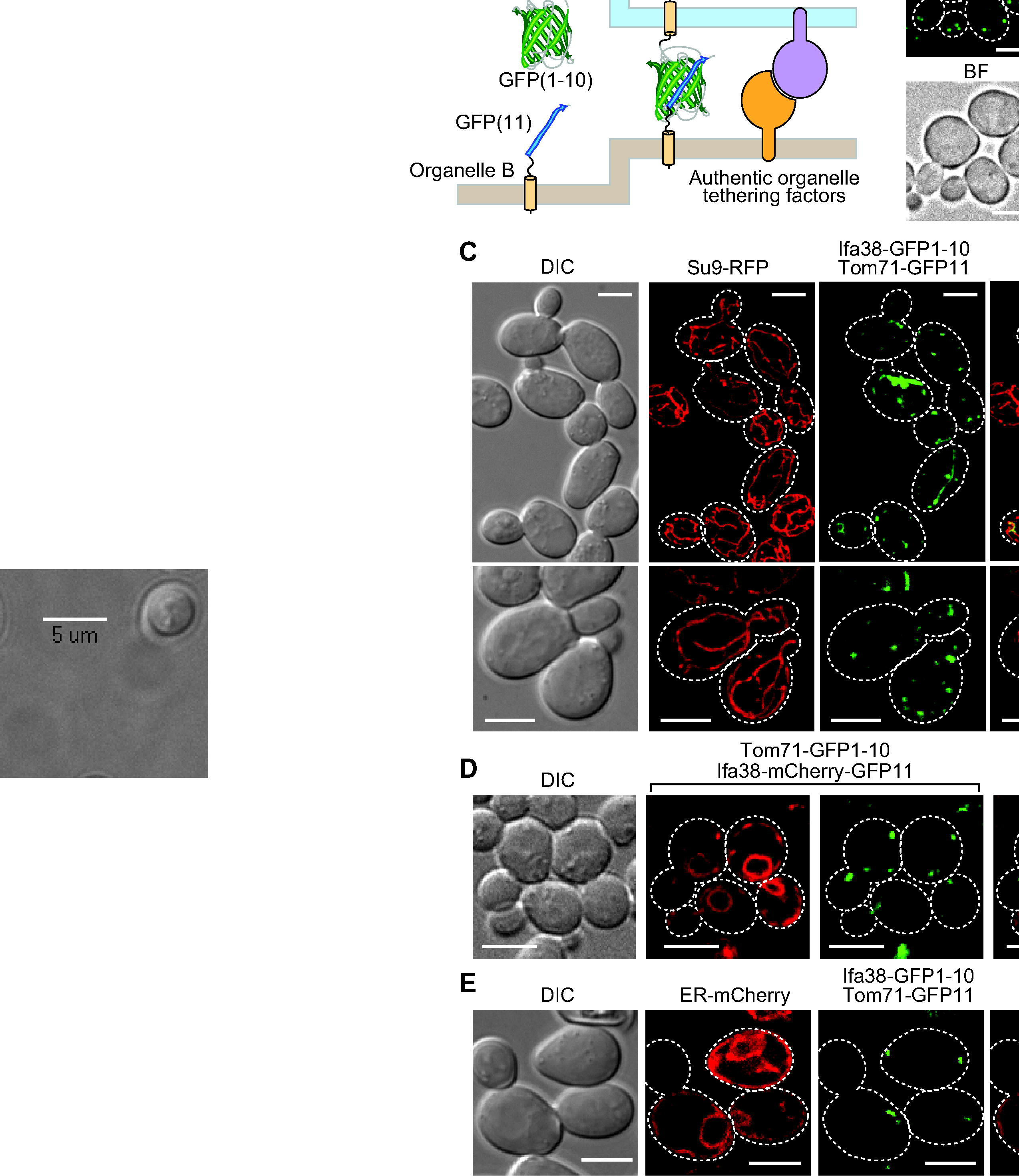
Split-GFP fragments on the ER and MOM label their contact sites. (**A**) Diagram of the split-GFP system for detecting inter-organelle interactions. (**B**) Yeast cells expressing Mmm1-GFP and mitochondria-targeted RFP (Su9-RFP) were imaged by confocal fluorescent microscopy. Maximum projection images reconstituted from the z-stacks were shown. Scale bar represents 5 µm. (**C**) Yeast cells expressing Tom71-GFP1-10 and Ifa38-mCherry-GFP11 were imaged by confocal fluorescent microscopy. Scale bar represents 5 µm. (**D, E**) Yeast cells expressing split-GFP proteins, Ifa38-GFP1-10 and Tom71-GFP11, and mitochondria-targeted RFP (Su9-RFP) (D) or the ER-targeted mCherry (E) were imaged by confocal fluorescent microscopy. Maximum projection (D) or a single focal plane images (D) were shown. Dotted lines indicate plasma membranes. Scale bar represents 5 µm.

We expressed split-GFP as fusion proteins with an ER membrane protein Ifa38 or a MOM protein Tom71, which are termed Ifa38-GFP1-10 and Tom71-GFP11, respectively. Since Ifa38 and Tom71 are anchored to the ER membrane and MOM with their N-terminal transmembrane (TM) segments from the cytosol, respectively, the split-GFP fragments fused to their C-terminus should be exposed to the cytosol. As a reference of authentic inter-organelle contact sites, we chose ERMES between the ER and mitochondria in yeast because ERMES is well characterized and can be easily observed as discrete foci under a fluorescent microscope by attaching a fluorescent-protein to Mmm1, one of the ERMES subunits (Fig 2B). We found that GFP signals generated from Ifa38-GFP1-10 and Tom71-GFP11 showed punctate structures (Fig 2C), which resemble the foci of the reference contact sites of ERMES (Fig 2B), although these fusion proteins are, like Ifa38 or Tom71 alone, expected to be uniformly distributed on each organelle as far as they do not interact with each other. We indeed found that, when Ifa38-mCherry-GFP11 and Tom71-GFP1-10 were co-expressed, the entire ER structure was stained with the mCherry signal from Ifa38-mCherry-GFP11, yet only specific parts of the ER were labeled with GFP signals from assembled Ifa38-mCherry-GFP11 and Tom71-GFP1-10 (Fig 2D). Besides, we confirmed that the GFP signals are present on mitochondria and the ER simultaneously, which are labeled with mitochondria-targeted RFP (Su9-RFP) and ER-targeted mCherry (ER-mCherry) (Fig 1C and E), respectively. This strongly suggests that the split-GFP fragments assemble at the ER-mitochondria contact sites. Essentially the same results were obtained when we used an ER protein Ynr021wp and N-terminal 33 residues containing the TM segments of human Tom20 as the ER-targeting and MOM-targeting signals, respectively (Fig. S1). We observed tubular GFP signals in rare cases (~10% of the cells), in which extensive, likely irreversible assembly of split-GFP fragments could expand the pre-existing contact-site regions (Fig 2C). Nevertheless, more than 90% of the cells exhibited only granular GFP signals, indicating that the effects of contact-site extension by the split-GFP expression remained minor.

### Assembled split-GFP signals are co-localized with pre-existing organelle contact sites

The granular GFP signals from Ifa38-GFP1-10 and Tom71-GFP11 indicate that the split-GFP system can be used to visualize ER-mitochondria contact sites efficiently. However, it is still possible that split-GFP fragments could spontaneously assemble on their own, independently of the pre-existing organelle contact sites, and that resulting irreversible association of the GFP fragments could actively promote formation of artificial ER-mitochondria contact sites. We thus examined if the GFP signals from the assembled split GFP fragments are co-localized with the ERMES foci. ERMES was stained with expressed Mdm34-RFP or Mdm12-mScarlet^32^. Strikingly, nearly all (88%) the GFP signals arising from assembled Ifa38-GFP1-10 and Tom71-GFP11 or assembled Ifa38-FLAG-GFP11 and Tom71-GFP1-10 were overlapped with the ERMES foci signals (Fig 3A and B). Furthermore, we confirmed that 82% of ERMES dots was stained with the split-GFP system. These observations clearly indicate that the split-GFP system is capable of labeling the pre-existing ER-mitochondria contact sites. These results also suggest that ERMES represents primary ER-mitochondria contact sites and that assembly of split-GFP fragments mainly takes place at the pre-existing ERMES regions on both organelles. Supporting this notion, expressions of the split-GFP fragments between mitochondria and the ER or vacuole hardly restored growth defects of ERMES-lacking *mmm1*∆ cells. This is in contrast to the cases of expression of ChiMERA, an artificial tethering protein between the ER and mitochondria, which was reported to partially rescue the growth defects^1^, and expression of the dominant-positive Vps13 mutant^11^ or Mcp1, both of which would induce mitochondria-vacuole contacts^12^ and indeed, at least partly relieved the growth defects (Fig 3C).

**Figure 3:**
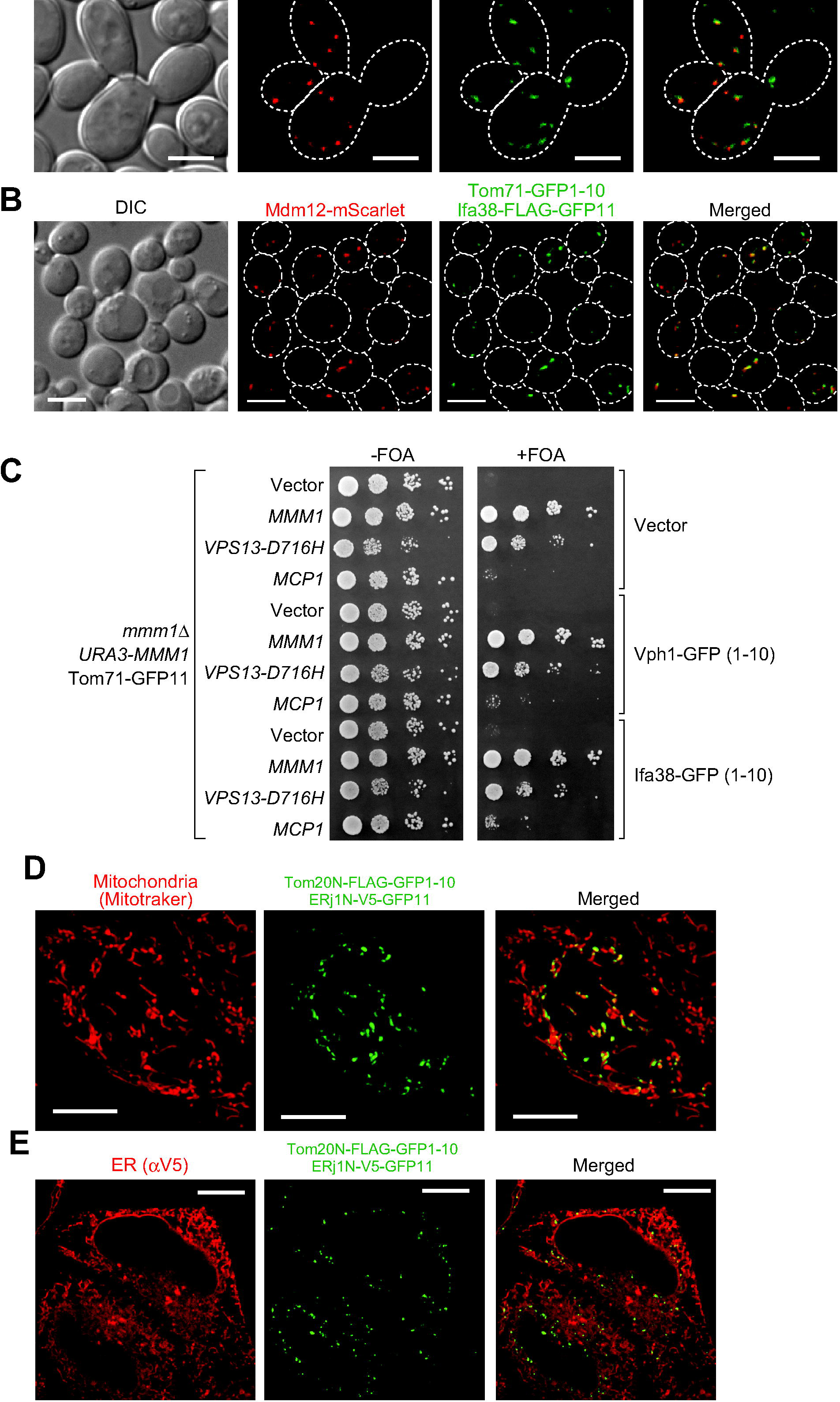
Co-localizations of assembled split-GFP fragments on the ER and mitochondria with ERMES. (**A, B**) Yeast cells expressing Mdm34-RFP from the multi-copy plasmid (pMY3^3^) (A) or Mdm12-mScarlet from the chromosome (B) were transformed with plasmids coding for the indicated split-GFP proteins and imaged by confocal fluorescent microscopy. Maximum projection images reconstituted from z-stacks were shown. Dotted lines indicate plasma membranes. Scale bar represents 5 µm. (**C**) *mmm1*∆ cells harboring a *CEN-URA3*-plasmid expressing Mmm1 (pYC1) were transformed with plasmids expressing the indicated split-GFP fusion proteins and Vps13-D716H, Mcp1, or empty vectors. Serial 10-fold dilutions of the resulting transformant cells were spotted onto selective synthetic media plates with or without 5’-FOA (5’-fluoroorotic acid). (**D**) HeLa cells transiently expressing Tom20N-FLAG-GFP1-10 and ERj1N-V5-GFP11 were stained with MitoTracker. Scale bar represents 10 µm. (**E**) HeLa cells used in (D) were subjected to immunofluorescence with anti-V5 antibodies. Scale bar represents 10 µm.

We then asked whether the split-GFP system can be also applied to detection of ER-mitochondria contact sites in mammalian cells by expressing split-GFP fragments on the ER membrane and MOM in HeLa cells. We expressed fusion proteins, the N-terminal 33 residues of a human MOM protein Tom20 followed by the FLAG tag and the split-GFP fragment (Tom20(1-33)-FLAG-GFP1-10) and the N-terminal 200 residues of an ER protein ERj1 followed by the V5 tag and the split-GFP fragment (ERj1(1-200)-V5-GFP11). Our live-cell imaging showed that these expressed split-GFP fusion proteins marked a part of mitochondria stained with MitoTracker in HeLa cells (Fig. 3D), which is essentially the same as what we observed for yeast cells (Fig. 2). We also confirmed that ERj1(1-200)-V5-GFP11 was distributed evenly on the ER by immunofluorescence using anti-V5 antibodies (Fig. 3E). Similar results were obtained when we used N-terminal 240 residues of human Sec63 and N-terminal 70 residues of human Tom70 as the ER- and MOM-targeting signals, respectively (Fig. S2). Taken together, we conclude that the split-GFP system could be applied to visualize inter-organelle contact sites in both yeast and mammalian cells.

### The Split-GFP fragments expressed on the ER and MOM can assemble in the absence of ERMES

In yeast, two different sets of tethering factors facilitating the ER-mitochondrial contact site formation were reported: the ERMES complex proteins and the EMC proteins plus Tom5^1,8^. Simultaneous deletion of the five EMC proteins, Emc1, 2, 3, 5 and/or 6 (5x-emc) does not cause decrease in the number of the ERMES foci, indicating that EMC is not essential for ER-mitochondria contact site formation^8^. However, it has not yet been tested whether deletion of the ERMES subunits has in turn a negative impact on ER-mitochondria contact-site formation although ERMES foci are completely gone when one of the ERMES subunits is depleted. This is due to lack of appropriate means to directly measure inter-organelle interactions irrespective of the presence of ERMES.

We thus tested the effects of loss of the ERMES subunits on formation of ER-mitochondria contact sites in yeast cells by using the split-GFP system. Interestingly, we observed that most *mmm1*∆ cells exhibited punctate GFP signals from Ifa38-GFP1-10 and Tom71-GFP11 (Fig. 4A) although a small population (~15%) of *mmm1*∆ cells contained spherical GFP signals, which outlined mitochondria with aberrant ball-like shapes (Fig. 4B). The spherical GFP signal may reflect the artificially formed ER-mitochondria contact sites on ball-shaped mitochondria, which are specific to *mmm1*Δ cells, like the tubular GFP signals observed in wild-type cells (Fig. 2C).

**Figure 4:**
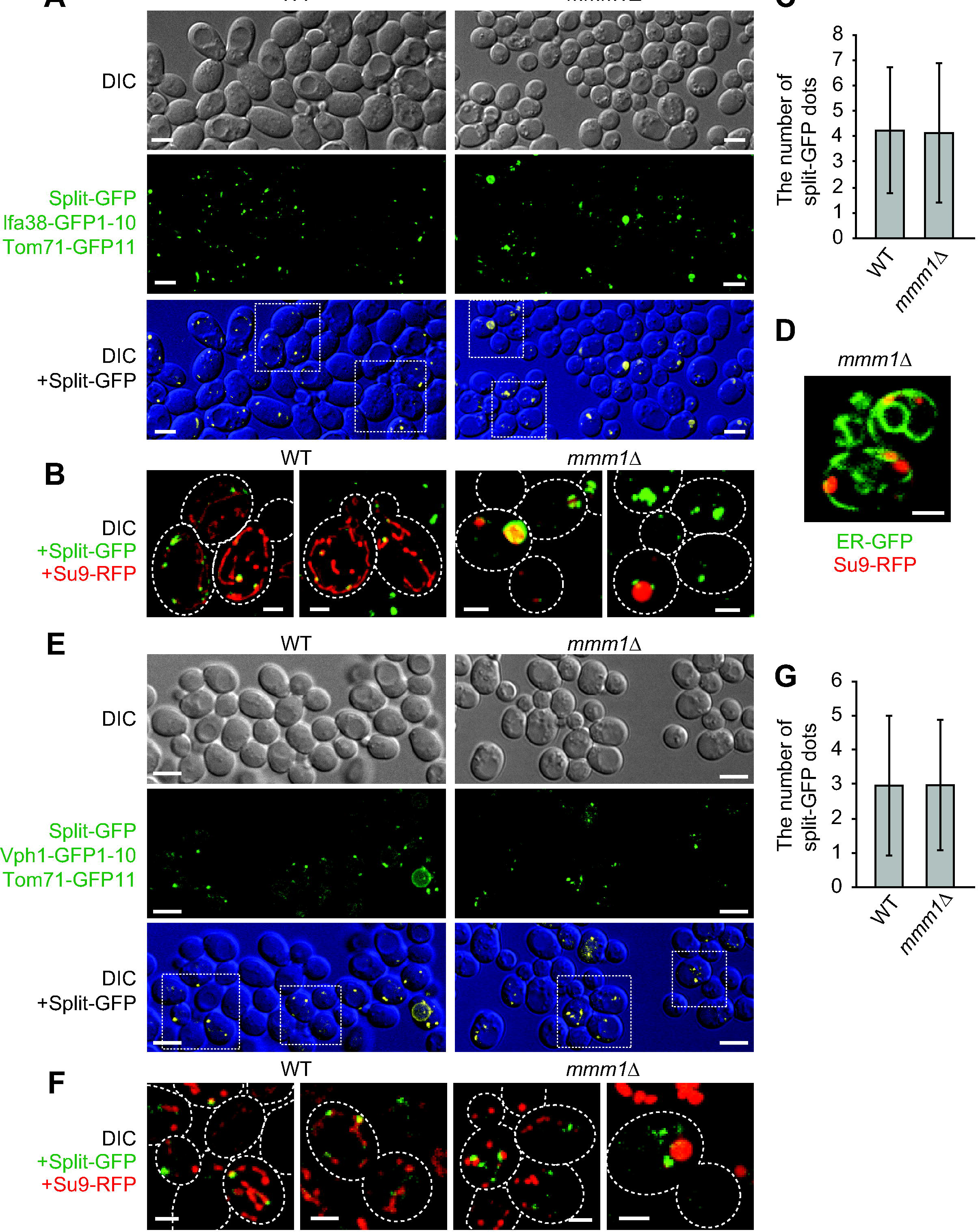
Loss of ERMES does not lead to a decrease in the number of split-GFP signals between the ER and mitochondria. (**A**) Wild-type and *mmm1*∆ cells expressing Ifa38-GFP1-10, Tom71-GFP11 and Su9-RFP were imaged by confocal fluorescent microscopy. Maximum projections reconstituted from z-stack GFP images were shown (middle panel) or overlaid with DIC images (lower panel). Scale bar represents 5 µm. (**B**) The maximum projection images surrounded by a rectangular frame in (A) were expanded and shown as merged images with su9-RFP. Dotted lines indicate plasma membranes. Scale bar represents 2 µm. (**C**) The number of GFP dot signals was counted. Error bars represent standard deviation. *n* = 87 and 115 for wild-type and *mmm1*∆ cells, respectively. (**D**) Yeast cells expressing ER-targeted GFP were stained with MitoTracker Red CMXRos and imaged by confocal fluorescent microscopy. A single focal plane image was shown. Scale bar represents 2 µm. (**E**) Wild-type and *mmm1*∆ cells expressing Vph1-GFP1-10, Tom71-GFP11, and Su9-RFP were imaged by confocal fluorescent microscopy. Maximum projections reconstituted from z-stack GFP images were shown (middle panel) or overlaid with DIC images (lower panel). Scale bar represents 5 µm. (**F**) The maximum projection images surrounded by a rectangular frame in (E) were expanded and shown as merged images with su9-RFP. Dotted lines indicate plasma membranes. Scale bar represents 2 µm. (**G**) The number of GFP dot signals was counted. Error bars represent standard deviation. *n* = 87 and 115 for wild-type and *mmm1*∆ cells, respectively.

To evaluate the contribution of ERMES to the ER-mitochondria contact-site formation, we compared the number of granular GFP signals per cell between wild-type and *mmm1*∆ cells. To count the number, we picked cells that exhibited only granular GFP signals. Surprisingly, *mmm1*∆ cells contained granular GFP signals whose number is comparable with those for wild-type cells (Fig. 4C). This suggest a possibility that ERMES is not essential for formations or maintenance of the ER-mitochondrial contact sites, and in the absence of ERMES, new contact sites can be formed by the factors other than ERMES subunits. In fact, we observed clear associations of mitochondria and the ER in the absence of Mmm1 by confocal microscopic analysis (Fig 4D). The split-GFP system thus provides a useful system to screen for novel factors that facilitate formation of mitochondria-ER contact sites.

The vCLAMP (mitochondria-vacuole contact) region was reported to drastically enlarge when ERMES is deficient^9^. We thus examined whether the Mmm1 loss affects the mitochondria-vacuole contact sites using the split-GFP system. To assess the contacts between mitochondria and the vacuole, we co-expressed Tom71-GFP11 and Vph1-GFP1-10, a fusion protein comprising of vacuolar membrane protein Vph1 followed by GFP1-10. Interestingly, we observed dot-like GFP signals from assembled Vph1-GFP1-10 and Tom71-GFP11 (Fig. 4E), like the ER-mitochondria contact sites visualized by Ifa38-GFP1-10 and Tom71-GFP11 (Fig 4A). These GFP signals were present on mitochondria (Fig. 4F) as well as on the vacuole (Fig. 5B), suggesting that the split-GFP fragments assembled between mitochondria and the vacuole. As shown in Fig. 4E, we did not see a drastic change in the overall pattern of the GFP signals from Vph1-GFP1-10 and Tom71-GFP11 between wild-type and *mmm1*∆ cells. The number and size of granular GFP signals were comparable between wild-type and *mmm1*∆ cells (Fig. 4E and G), indicating that loss of ERMES does not lead to increase in the number of mitochondria-vacuole contact sites.

**Figure 5:**
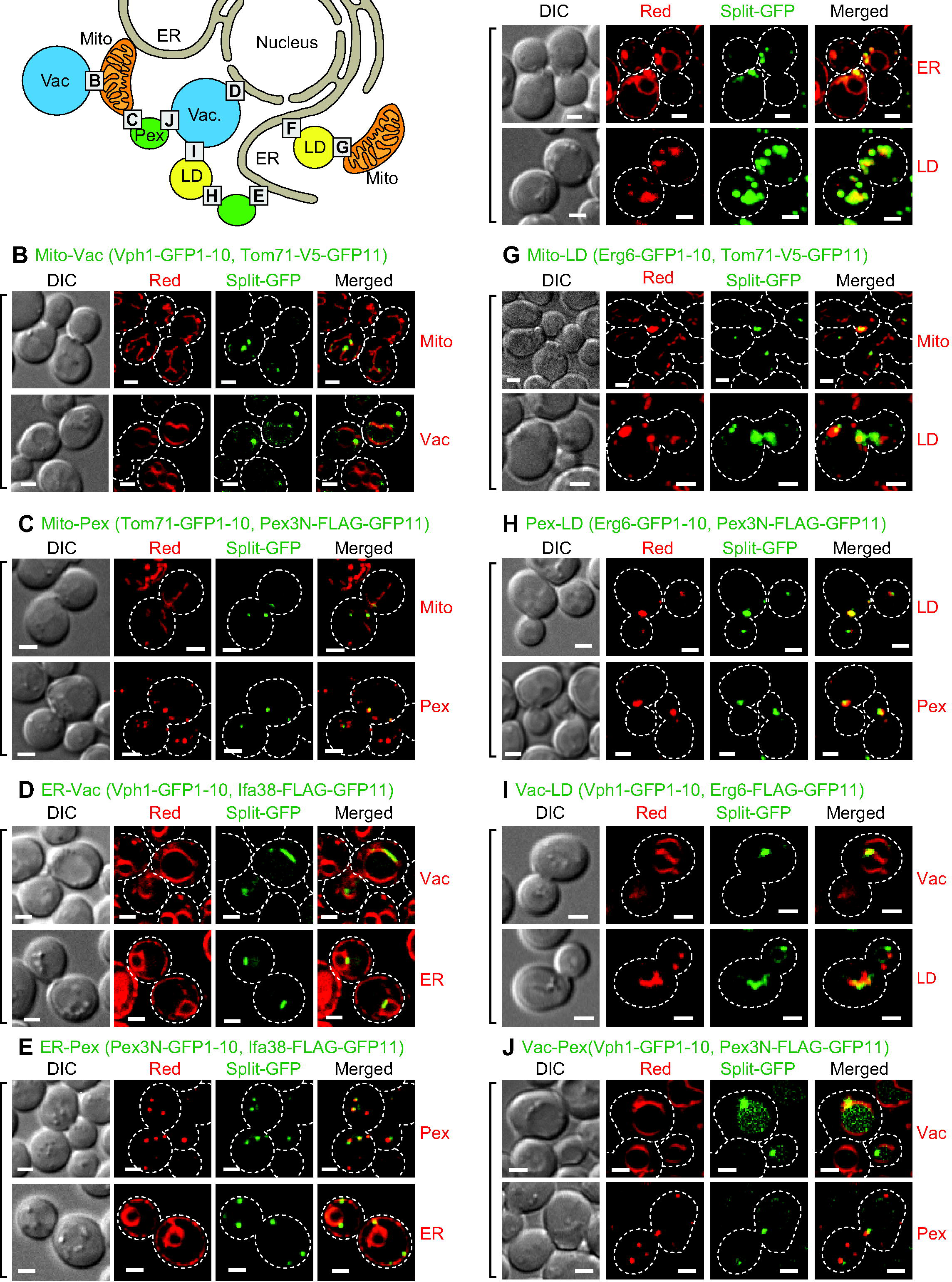
A number of inter-organelle contact sites visualized with the split-GFP system. (**A**) Diagram of inter-organelle contact sites we tested. The alphabet X in squares correspond to Fig 4X showing the corresponding inter-organelle contact sites. (**B-J**) Wild-type cells expressing the indicated split-GFP proteins were imaged by confocal fluorescent microscopy. Su9-RFP, mCherry-PTS1, BipN-mCherry-HDEL and Erg6-mCherry were used to label mitochondria, peroxisomes, the ER and LDs, respectively (organelle marker proteins were summarized in Materials and Methods). The vacuole and LDs were stained with fluorescent dyes, FM4-64 and BODIPY 558/568 C_12_, respectively. Maximum projection was performed to show split-GFP signals on mitochondria, peroxisomes and LDs while a single focal plane was shown for the ER and vacuole. Dotted lines indicate plasma membranes. Scale bar represents 2 µm.

### The split-GFP system can be used to detect various inter-organelle contact sites

In addition to the ER-mitochondria and mitochondria-vacuole pairs, several different organelle pairs were shown to be in close proximity in yeast cells^17–20,29^. Nevertheless, proteins responsible for formation of those possible organelle contact sites largely remain to be identified (Fig. 5A). We thus asked whether the split-GFP system could be used to detect not only ER-mitochondria and mitochondria-vacuole contact sites but also other inter-organelle contact sites. To anchor split-GFP fragments to LD and peroxisomal membranes from the cytosol, we utilized a LD protein Erg6, and the N-terminal 60 residues of a peroxisomal membrane protein Pex3 containing a TM segment (Pex3N) (Fig. 1). By using these differently targeted split-GFP constructs, we tested all combinations of the organelle pairs among the ER, mitochondria, vacuole, peroxisomes and LDs for contact-site observations.

Strikingly, we were able to observe clear punctate GFP signals for all the organelle pairs we tested, like the case of the ER-mitochondria and mitochondria-vacuole contact sites (Fig. 5B–J). Since these split-GFP-fusion proteins should be evenly distributed on the entire target organelles, the punctate GFP signals strongly suggest that the split-GFP fragments assemble and accumulate at the organelle contact sites where two distinct organelles are juxtaposed. In particular, the split-GFP signals from vacuole-targeted Vph1-GFP1-10 and ER-targeted Ifa38-FLAG-GFP11 were observed only around the nuclear ER but not the peripheral ER while Ifa38 is localized in both the nuclear and peripheral ER (Figs. 5D and 2D). We also noticed that the split-GFP signals from the ER and vacuole pair appeared as short tubules rather than dots (Fig 5D). This short tubular shape is similar to that of the authentic NVJ signal reported previously^21–25^. Therefore these observations also support the idea that the split-GFP fragments do not strongly induce organelle contact sites. In addition, our results suggest that NVJ (nuclear ER) is the major site where the ER and vacuole interact with each other. It is also to be noted that the vacuole-peroxisome contact sites (Fig. 5J) were not previously reported although organelle contact sites between peroxisomes and organelles other than the vacuole were reported^15,16,28,29,33^. Taken together, we conclude that the split-GFP system is potentially useful for efficient detection of contact sites between any organelle membranes in living cells.

## Discussion

While inter-organelle contacts are recognized as important structures for coordinated functions of different organelles, detailed analyses of such contact sites were hampered due to the lack of a reliable system to observe the regions or contact sites where various pairs of distinct organelle membranes are closely apposed. A widely used method to detect inter-organelle contact sites employs a fluorescent protein attached to a known component of the contact sites such as ERMES and the NVJ, but then it is naturally difficult to observe or analyze the contact sites in the absence of ERMES of the NVJ. In the present study, we evaluated the usability of the split-GFP system to visualize organelle contact sites in yeast and mammalian cells by live-cell imaging. Importantly, our results strongly suggest that expression of the split-GFP fragments targeted to different organelle membranes may not artificially induce new contact sites between organelles due to the following reasons. First, most GFP signals arising from the split-GFP system showed granular signals rather than tubular or aggregated patterns. Second, expression of split-GFP fragments targeted to the ER and mitochondrial membranes provides GFP signals co-localized with the ERMES dots, which represented the pre-existing authentic ER-mitochondria contact sites (Fig. 3A and B). Third, the expression of split-GFP fragments targeted to the ER and mitochondrial membranes did not substantially rescue the growth defects of *mmm1*∆ cells (Fig. 3C), while such growth defects were partially relieved by expressed ChiMERA, a nonfunctional artificial tether between the ER and mitochondria^1^, and dominant-positive Vps13 mutant^11^ or Mcp1, both of which would induce mitochondria-vacuole contacts^12^. Forth, split-GFP probes targeted to the ER and vacuole membranes labeled exclusively the nuclear ER region that was closely apposed to the vacuole (Fig. 5D), which resembled the NVJs reported previously^21–25^. However, we cannot rule out the possibility that the split-GFP system perturbs the physiological organelle-contact sites in some cases. For example, we noticed that mitochondrial shape looked partly aberrant when split-GFP fragments were expressed between mitochondria and peroxisomes or between mitochondria and LDs; although yeast cells normally show elongated tubular mitochondria (Fig. 2B and C), mitochondrial tubules became rather short when split-GFP fragments were expressed on the MOM and peroxisomes or LDs (Fig. 5C and G). These observations may indicate that the split GFP system could affect the organelle morphology or functions by perturbing the inter-organelle contacts to some extent.

Interestingly, the mitochondria-ER contact sites visualized by the split-GFP system do not simply fit the current understanding of the role of ERMES as primary ER-mitochondria contact sites; the number of dots from the assembled split GFP signals did not change with and without Mmm1 of the ERMES complex. One likely explanation for this is that unidentified ER-mitochondria tethering factors may be up-regulated to compensate the loss of ERMES. In fact, our microarray analyses using total mRNAs showed that mRNA levels of a considerable number of genes including several uncharacterized ones were significantly increased in *mmm1*∆*mdm12*∆ cells as compared with those in wild-type cells (data not shown). Identification of such additional tethering factors between the ER and mitochondria should be an essential subject in future studies.

Another observation that is not consistent with the previously reported one is concerned with the role of vCLAMP in formation of the mitochondria-vacuole contact sites. While the vCLAMP region was reported to enlarge significantly upon loss of an ERMES component^9^, we did not observed such increases in the mitochondria-vacuole contact-site sizes or numbers with the split-GFP system (Fig. 4E and G). However, we noted that even in the previous study, expansion of vCLAMP was observed for a part of the cells, but not all the cells^9^. This could raise alternative possible interpretation that variation in the expression level of GFP-Vps39, but not the loss of the ERMES component, may be the cause for the enlarged vCLAMP observed in the previous study^9^. Supporting this idea, a recent study showed that overexpression of Vps39 simply expanded the vCLAMP region and that chromosomally expressed Vps39-GFP exhibited dot-like patterns^12^ similar to those observed with our split-GFP system (Fig. 4E and F). Therefore, the expansion of vCLAMP in response to the loss of ERMES is still controversial, and it remains be addressed which of the systems is more suitable for visualizing the intact vCLAMP regions, the split GFP system or Vps39-GFP.

Our systematic observations for various pairs of organelles among the ER, mitochondria, vacuole, peroxisomes, and LDs strongly suggest that each organelle simultaneously forms multiple contact sites with various organelles (Fig. 5). Although tethering factors were identified for ERMES and the NVJ, factors for forming other inter-organelle contact sites, including those between LDs and peroxisomes identified here, still remain obscure. Since the present split GFP system allows us to evaluate the presence of inter-organelle contacts with minimum cellular perturbations, it can be used for systematic screening for the factors responsible for formation and regulation of inter-organelle contact sites. Uncovering the physiological roles of each organelle contact site and organelle-tethering factor will be the next important issues we have to tackle.

## Materials and Methods

### Yeast strains and growth conditions

FY833 was used as wild-type strain throughout this research^34^. Yeast cells expressing Mmm1-GFP (Mmm1-GFP) or those lacking Mmm1 (*mmm1*∆) were described previously^35^. The mScarlet tag^32^ was inserted proximal to the stop codon of *MMM1* by homologous recombination using gene cassettes from pFA6a-mScarlet-KanMX (see below). Cells were grown in SD (0.67% yeast nitrogen base without amino acids, 0.13% drop-out amino acid mix and 2% glucose) or SCD (0.67% yeast nitrogen base without amino acids, 0.5% casamino acids and 2% glucose), both of which were further supplemented appropriately with 20 µg/ml each of adenine, L-tryptophan, L-histidine, L-methionine, and uracil and 30 µg/ml each L-leucine and L-lysine. The drop-out amino acid mix was the mixture of 2.6 g adenine, 6.0 g L-aspartic acid, 12 g L-threonine, 2.6 g L-asparagine, 1.8 g L-tyrosine, 6.0 g L-glutamic acid, 2.6 g L-glutamine, 2.6 g glycine, 2.6 g L-alanine, 2.6 g L-isoleucine, 1.2 g L-methionine, 3.0 g L-phenylalanine, 2.6 g L-proline, 22.6 g L-serine, 9.0 g L-valine and 2.6 g L-cysteine.

### Plasmids

Plasmids and primers used in this study are summarized in Tables S1 and S2, respectively. For the expressions of split-GFP-fusion proteins or organelle marker proteins in yeast, we first prepared yeast expression vectors with the *GPD* or *ADH1* promoter and the *CYC1* terminator in *CEN*-plasmids, pRS313, pRS314, pRS315 and pRS316^36^, resulting in pYU41, pYU47, pYU53, pYU54 and pYU59. Then, synthesized DNA fragments coding for GFP1-10, GFP11, FLAG-GFP11 or V5-GFP11 (Supplementary information), purchased from Eurofins Genomics, were cloned into BamHI/EcoRI sites of pYU47 or pYU59, resulting in pSFL9, 10, 11, 12, 73 or 74 (Table S1). Then, DNA fragments coding *YNR021W*, *IFA38*, *VPH1*, *TOM71*, *PEX3(1-60)* or *ERG6* were amplified by PCR with pairs of primers, #YU291 and 292, #YU293 and 294, #YU295 and 296, #YU297 and 298,#YU305 and 306 and #YU307 and 308, respectively and cloned into NotI/BamHI sites of pSFL9, 10, 11, 12, 73 or 74. In some cases, tandem genes coding for an organelle protein and the split-GFP fragment were cut with NotI and EcoRI or HindIII, and then inserted into pYU41 and pYU53.

To visualize the ER, peroxisomes and LDs, fusion proteins named BipN-mCherry-HDEL, mCherry-PTS1 and Erg6-mCherry were expressed from the *CEN*-plasmids, pFL16, pFL24 and pFL72, respectively. The DNA fragments coding for N-terminal 47 residues of Kar2 (BipN), mCherry-HDEL (the ER retention signal), mCherry-PTS1 (peroxisome targeting signal 1, SKL), Erg6 and mCherry were amplified by PCR with pairs of primers, #NU892 and 893, #NU946 and 948, #YU377 and 378, #YU307 and 308, and #NU539 and 540, respectively. The resulting DNA fragments coding for BipN and mCherry-HDEL were inserted tandemly into pYU59, resulting in pFL16. The resulting DNA fragment coding for mCherry-PTS1 was inserted tandemly into pYU59, resulting in pFL24. The resulting DNA fragments coding for Erg6 and mCherry were inserted tandemly into pYU59, resulting in pFL72. For expression of Mdm34-RFP, plasmid pMY3 was used^3^.

To construct pFA6a-mScarlet-KanMX, a synthesized DNA fragment coding for mScarlet was purchased from Eurofins Genomics and inserted at PacI/PmeI sites in pFA6a-3HA-kanMX6^37^. pMM80, 82, 87 and 89, plasmids for expressions of split-GFP-fusion proteins in HeLa cells were constructed as follows. First, DNA fragments coding for Tom20 (1-33)-FLAG, Tom70(1-70)-FLAG, ERj1(1-200)-V5 or Sec63(1-240)-V5 were cut from pMM73 to 77, and inserted into BamHI-NotI-cut pcDNA3.1+C-eGFP. The resulting plasmids were then digested BamHI and XbaI and ligated with BamHI-XbaI-digested DNA fragments coding for GFP1-10 or GFP11 amplified by PCR using a pair of primers #YU1006 and 1008 or #YU1007 and 1009 and pSFL1 or pSFL2 as template (Table S1). Plasmids pMM73 to 77 were purchased from Genscript.

### Cell culture and transfection

HeLa cells were maintained at 37°C in DMEM supplemented with 10% FBS. DNA transfection was performed using Lipofectamine 2000 (Invitrogen) according to the manufacturer’s instructions. Briefly, 24 hours before transfection, HeLa cells were seeded in 35 mm glass-bottom dish (Iwaki) with a seeding density of 1.5 − 10^5^ in 2 ml DMEM supplemented with 10% FBS and incubated. Then, the HeLa cells were co-transfected with two plasmids for the expressions of GFP1-10- and GFP11-fusion proteins (1.25 µg each/ 35 mm dish) and further incubated for 24 hours for the microscopic analysis.

### Fluorescence microscopy

Yeast cells were grown in SCD or SD medium supplemented with appropriate amino acids to keep plasmids expressing the split-GFP-fusion proteins and organelle marker protein. Logarithmically growing cells were observed under Olympus IX83 microscope with a CSU-X1 confocal unit (Yokogawa), a 100 x, 1.4 NA, objective (UPlanSApo, Olympus) and an EM-CCD camera (Evolve 512; Photometrics) manipulated by Metamorph software (Molecular Devices). GFP or RFP, mCherry and cy5 were excited by 488-nm or 561-nm laser (OBIS, Coherent) and the emission was passed through 520/35-nm or 617/73-nm band-pass filter, respectively. The confocal fluorescent sections were collected every 0.2 µm from the upper to bottom surface of yeast cells. For spit-GFP images labeled with mitochondria, peroxisomes and LDs, the obtained confocal images were subjected to maximum projection using Image J software.

For immunofluorescence microscopy, HeLa cells grown in 35 mm glass-bottom dish were fixed for 15 min at room temperature with pre-warmed 4% PFA in phosphate buffer and washed three times with PBS. Cells were permeabilized with 0.5% Triton X100 in PBS for 15 minutes and washed three times with PBS. After blocking with 3% BSA containing PBS for 1 hour, cells were incubated with either 1 μg/ml anti-V5-tag mAb (MBL life science) in blocking buffer for 1 hour. Cells were washed three times with PBS and incubated with 2 μg/ml goat anti-mouse IgG (H+L) Cross-Adsorbed Secondary Antibody, Cyanine5, (Thermo Fisher Scientific) in blocking buffer for 1□hour. For MitoTracker staining, cells were incubated with 100 ng/ml MitoTracker Red CMXRos (Thermo Fisher Scientific) in Opti-MEM for 30 minutes at 37□°C in 5% CO_2_. Cells were washed two times with DMEM supplemented with 10% FBS and subjected to microscopic observation.

## Acknowledgements

We thank Dr. Hiromi Sesaki (Johns Hopkins University School of Medicine) for pYM3 and Su9-RFP plasmids, and K. Shishido for her great technical assistance. We are grateful to members of the Tamura and Endo laboratories for helpful discussion. This work was supported by JSPS KAKENHI (Grant Numbers 15H05595 and 17H06414 to YT, and 15H05705 and 22227003 to TE), Senri Life Science Foundation (YT) and a CREST Grant from JST (TE). ST is a Research Fellow of the JSPS.

## Author contributions

TE and YT designed the study. YK performed the most yeast experiments. ST performed the experiment using HeLa cells. RK performed the yeast growth assay and analyzed the number of GFP signals. YM was involved in the initial phase of this study. TE and YT wrote the paper.

## Additional Information

**Supplementary information** accompanies this paper at http://www.nature.com/srep

