## Supplementary Materials for "Visualizing multiple inter-organelle contact sites using the organelle-targeted split-GFP system" (from another version of this preprint: version 1, 2018-03-27)

**Contents**

**Supplementary figures S1 to S6, Materials and Table S1 and S2**

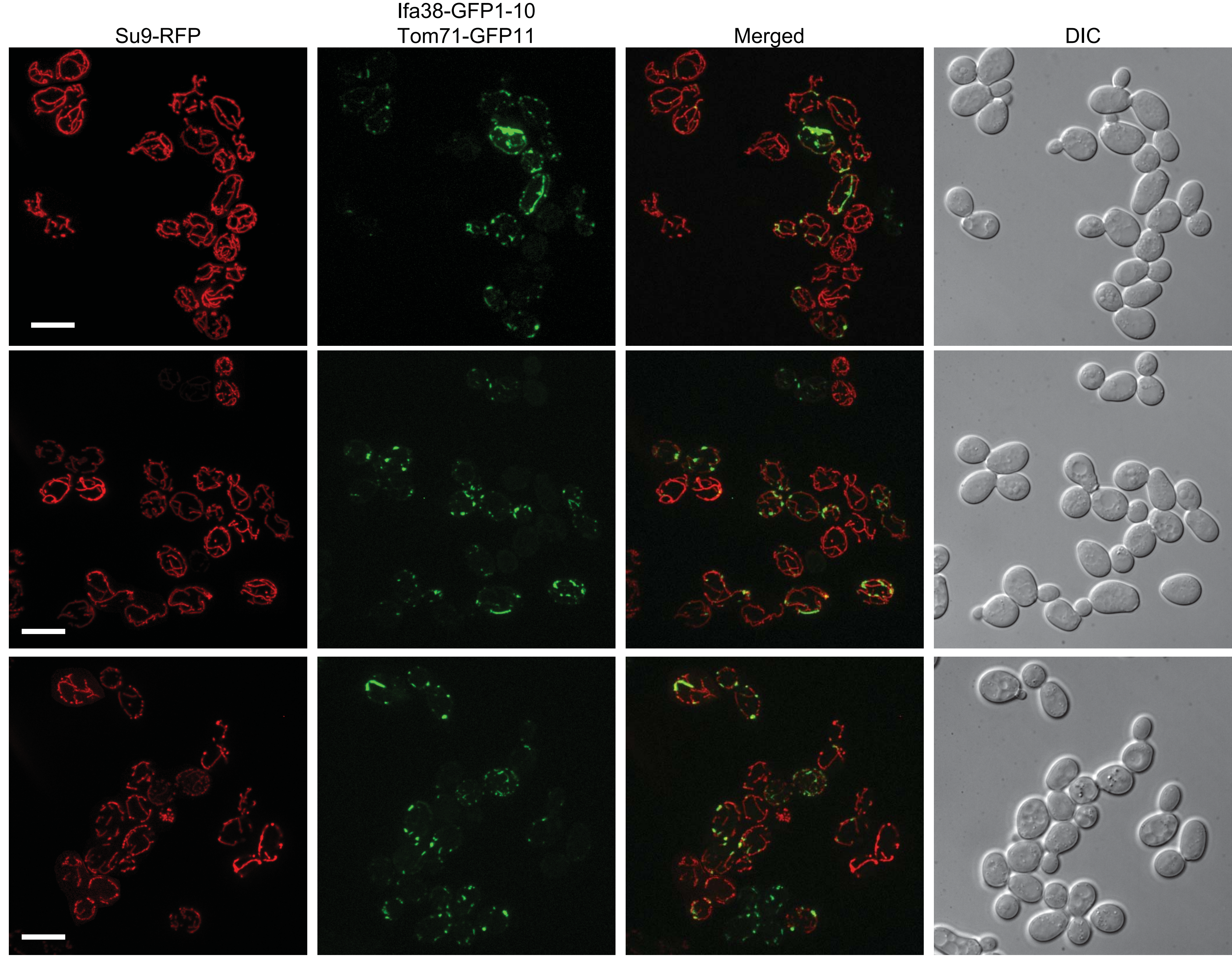


**kakimoto et al., Figure S1**

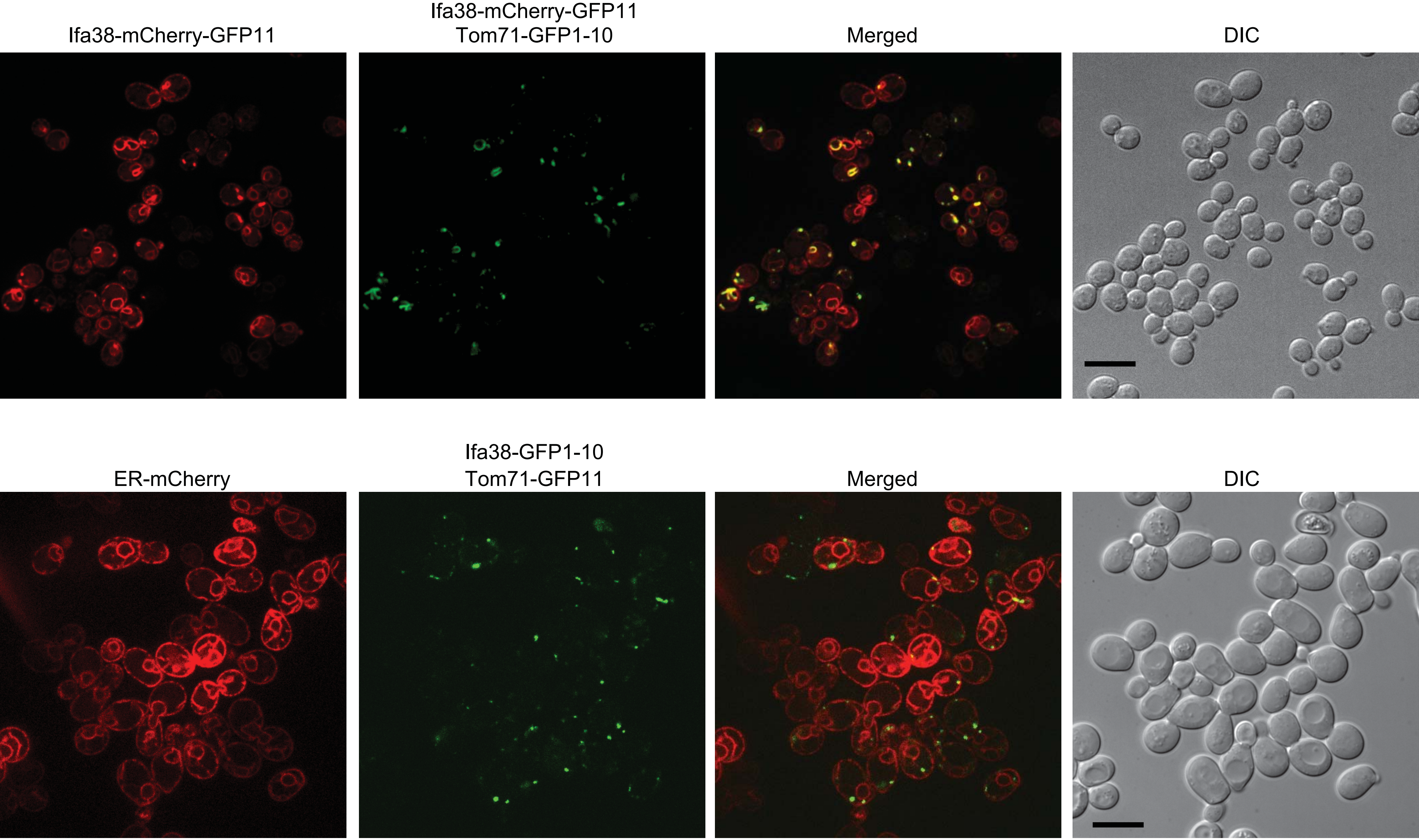


**kakimoto et al., Figure S2**

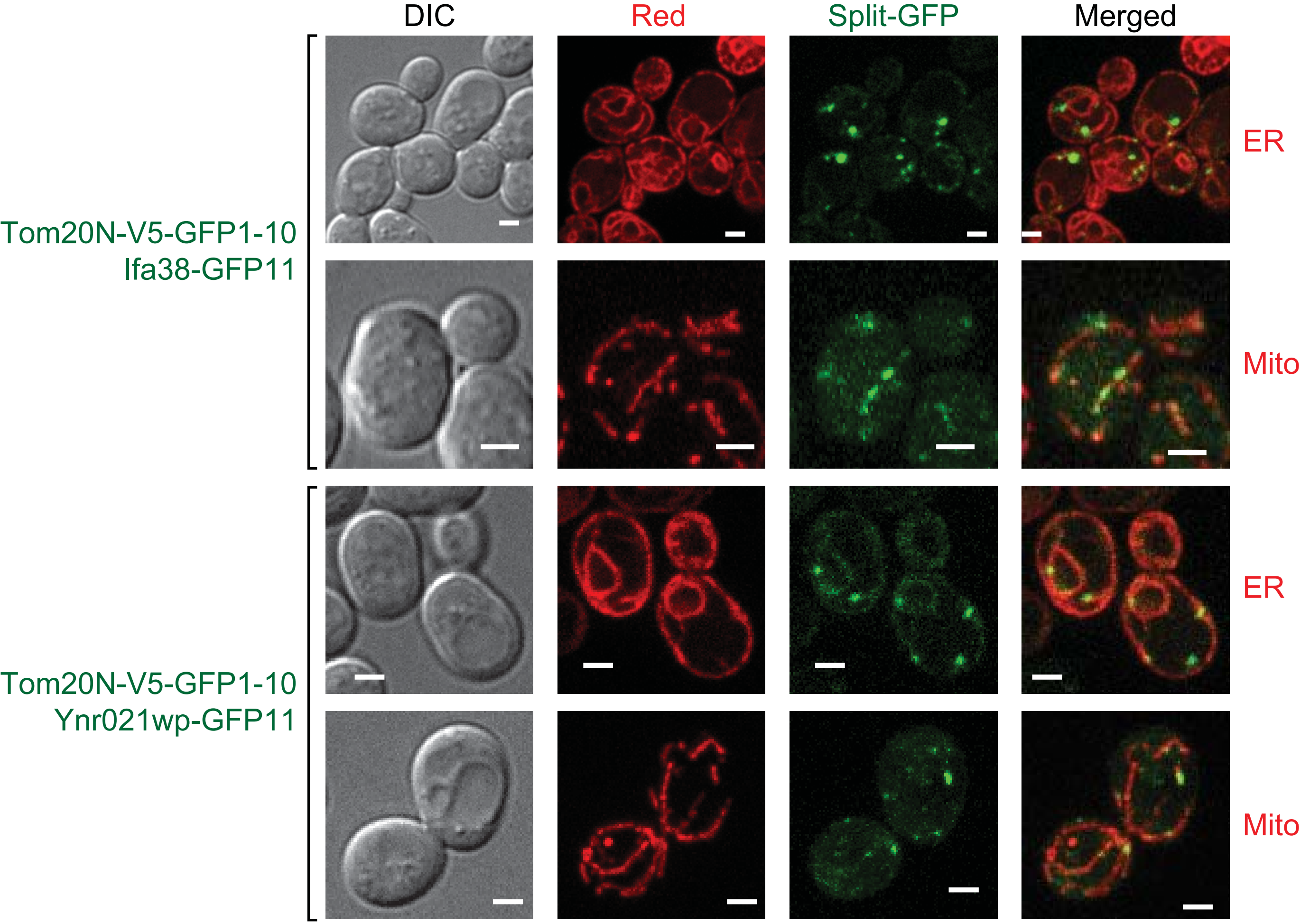


**kakimoto et al., Figure S3**

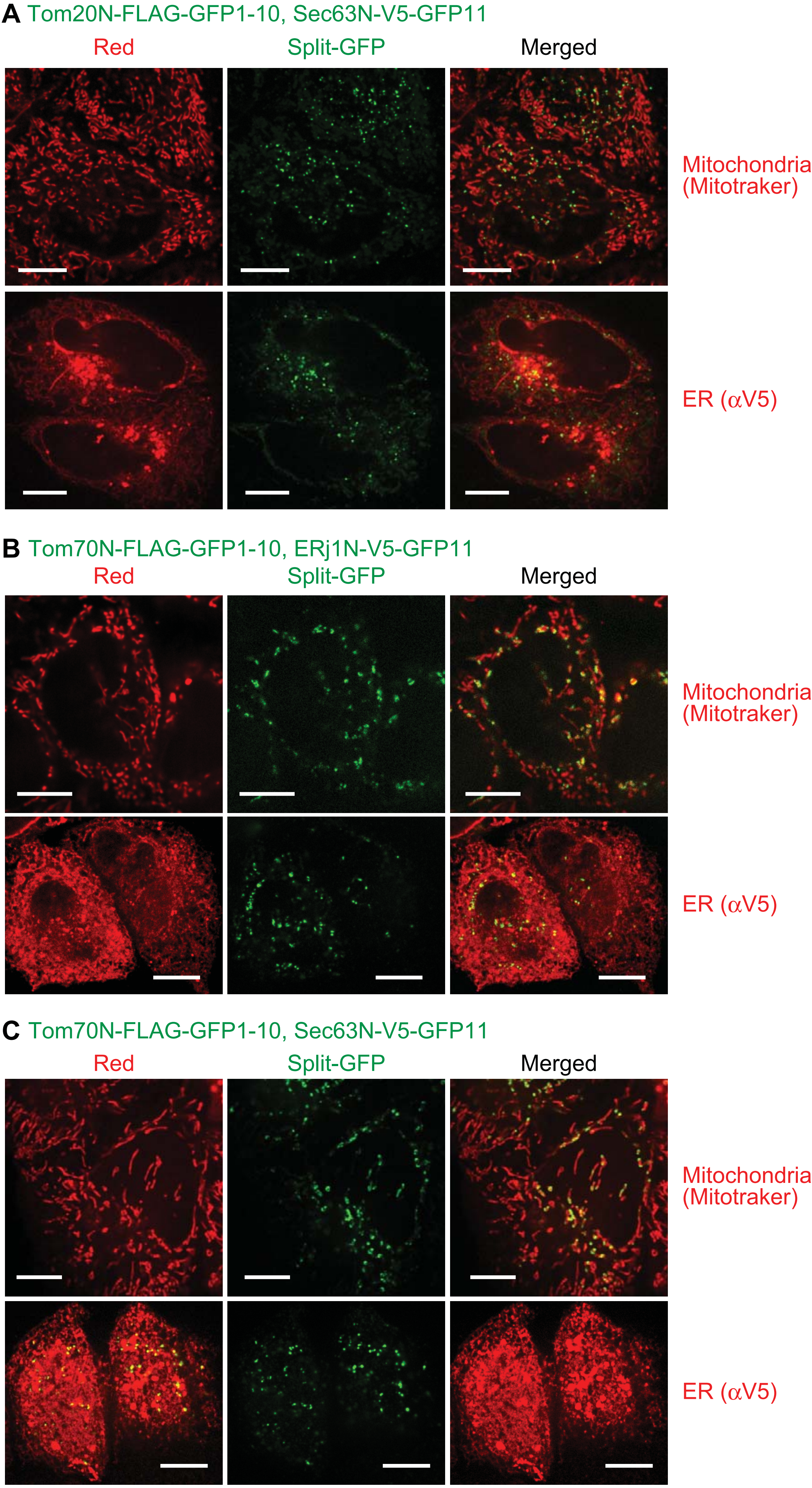


**kakimoto et al., Figure S4**

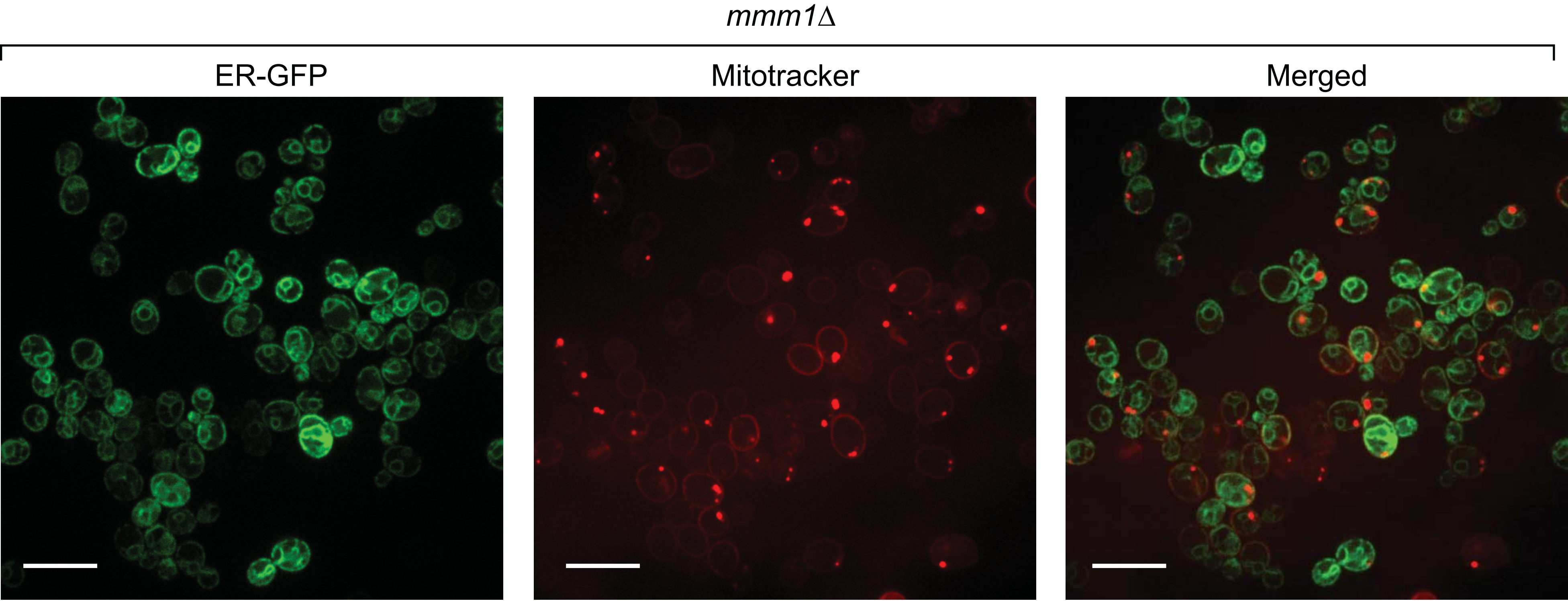


**kakimoto et al., Figure S5**

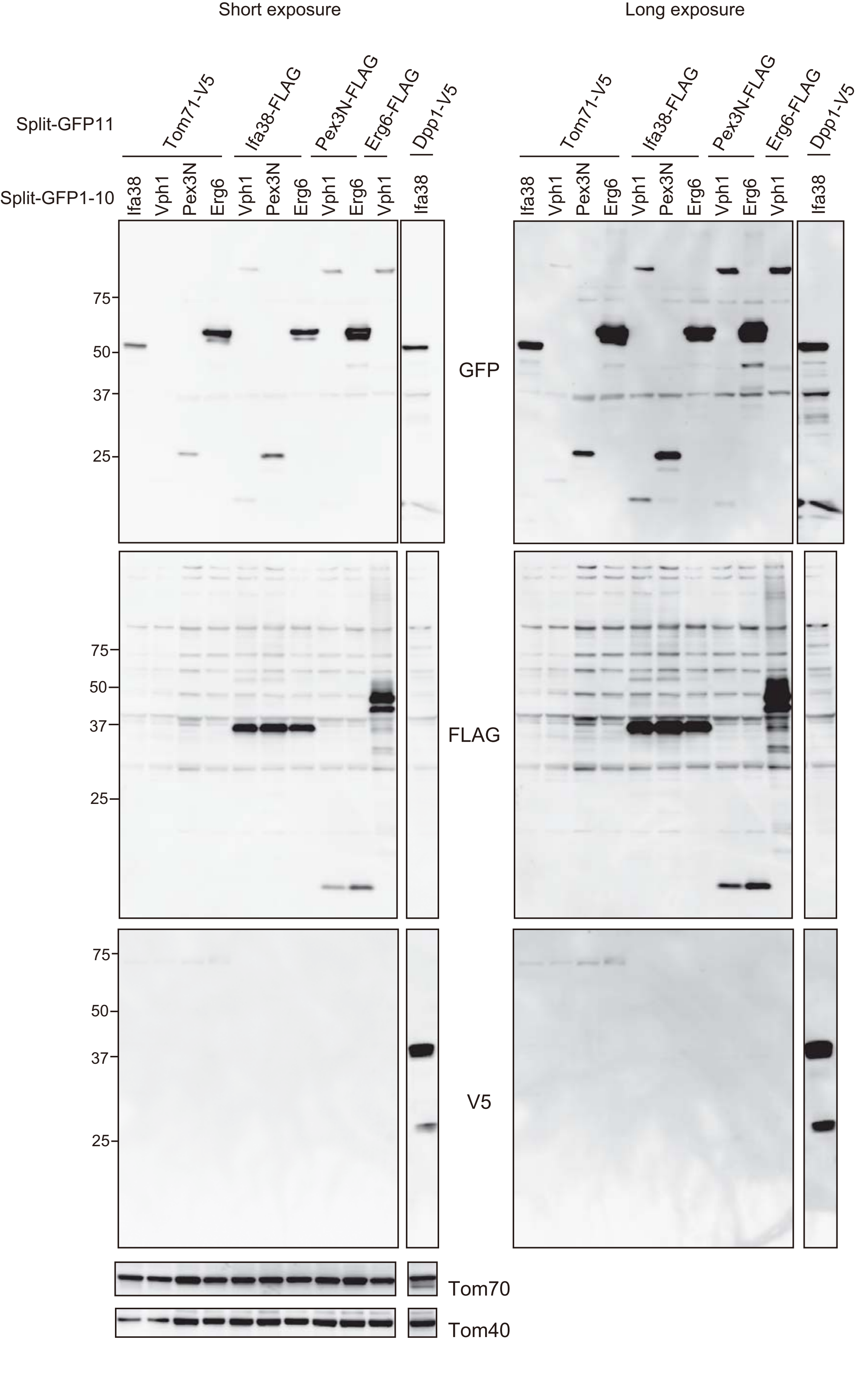


kakimoto et al., Figure S6

**Figures S1 and S2. Representative images showing cells expressing split-GFP probes for visualizing the ER-mitochondria contact sites.** Yeast cells expressing Ifa38-GFP1-10 and Tom71- GFP11 (S1), Ifa38-mCherry-GFP and Tom71-GFP1-10 (S2A) and Ifa38-GFP1-10 and Tom71-GFP11 (S2B) were imaged by confocal fluorescent microscopy. Maximum projection images of whole visual microscope field were reconstituted from the z-stacks and were shown. Scale bar represents 10 μm.

**Figure S3. The ER-mitochondria contact sites visualized with split-GFP fragments in yeast.**

Yeast cells expressing split-GFP proteins, Tom20N-V5-GFP1-10 and Ifa38-GFP11 or Ynr021wp-GFP11, and mitochondria-targeted RFP (Mito) or the ER-targeted mCherry (ER) were imaged by confocal fluorescent microscopy. Scale bar represents 2 μm.

**Figure S4. The ER-mitochondria contact sites visualized with the split-GFP system in HeLa cells.**

A-C HeLa cells transiently expressing Tom20N-FLAG-GFP1-10 and Sec63N-V5-GFP11 (A), Tom70N-FLAG-GFP1-10 and ERj1N-V5-GFP11 (B), Tom70N-FLAG-GFP1-10 and Sec63N-V5-GFP11 (C), were stained with MitoTracker and immediately observed (upper panel) or fixed and subjected to immunofluorescence using anti-V5 antibodies (lower panel). Scale bar represents 10 μm.

**Figure S5. The ER and mitochondria associate in the absence of Mmm1.**

Yeast cells expressing ER-targeted GFP were stained with MitoTracker Red CMXRos and imaged by confocal fluorescent microscopy. Single focal plane images of whole visual microscope field were shown. Scale bar represents 10 μm.

**Figure S6. Immunoblotting of total cell lysates showing the expression levels of split-GFP probes.** Total cell lysates were prepared from logarithmically growing cells expressing the indicated split-GFP fusion proteins and were analyzed by immunoblotting using the indicated antibodies. Arrowheads indicate degraded products from Vph1-GFP1-10.

### Materials

### DNA sequences for the split-GFP fragments and organelle-targeting signals

#### GFP1-10

CATGGGTGGCACTAGTATGAGCAAAGGAGAAGAACTTTTCACTGGAGTTGT
CCCAATTCTTGTTGAATTAGATGGTGATGTTAATGGGCACAAATTTTCTGTC
AGAGGAGAGGGTGAAGGTGATGCTACAATCGGAAAACTCACCCTTAAATTT
ATTTGCACTACTGGAAAACTACCTGTTCCATGGCCAACACTTGTCACTACTC
TGACCTATGGTGTTCAATGCTTTTCCCGTTATCCGGATCACATGAAAAGGCA
TGACTTTTTCAAGAGTGCCATGCCCGAAGGTTATGTACAGGAACGCACTATA
TCTTTCAAAGATGACGGGAAATACAAGACGCGTGCTGTAGTCAAGTTTGAA
GGTGATACCCTTGTTAATCGTATCGAGTTAAAGGGTACTGATTTTAAAGAAG
ATGGAAACATTCTCGGACACAAACTCGAGTACAACTTTAACTCACACAATG
TATACATCACGGCAGACAAACAAAAGAATGGAATCAAAGCTAACTTCACAG
TTCGCCACAACGTTGAAGATGGTTCCGTTCAACTAGCAGACCATTATCAACA
AAATACTCCAATTGGCGATGGCCCTGTCCTTTTACCAGACAACCATTACCTG
TCGACACAAACTGTCCTTTCGAAAGATCCCAACGAAAAGTAA

#### GFP11

CGATGGAGGGTCTGGTGGCGGATCAACAAGTCGTGACCACATGGTCCTTCA
TGAGTACGTAAATGCTGCTGGGATTACATAA

#### 3xFLAG-GFP11

TCTGCCGGTGGTGATTACAAAGACCATGATGGCGATTATAAGGATCATGAC
ATTGACTATAAGGATGATGACGATAAAGGCGGAAGTGGTGGAGGTTCAACT
TCCAGAGATCACATGGTTTTGCATGAATACGTCAATGCTGCAGGGATAACA
TAA

#### V5-GFP11

TCTGCCGGAGGTTCTGGGAAACCCATACCAAACCCTCTTTTGGGCTTAGACT
CAACTGGTGGAAGTGGTGGTGGTTCCACAAGCAGAGATCATATGGTTCTAC
ACGAATATGTCAATGCTGCAGGCATTACCTAA

**ERj1(1-200)-V5**

GCCACCATGACTGCTCCGTGCTCTCAACCCGCACAACTGCCAGGACGGAGA
CAGCTGGGTCTGGTACCTTTTCCTCCTCCACCACCACGTACACCTCTGCTGT
GGTTGCTGCTTCTGCTGCTTGCTGCGGTTGCACCGGCTAGAGGATGGGAATC
CGGCGATTTGGAGCTGTTTGACCTGGTCGAAGAAGTGCAGTTGAACTTCTAC
CAGTTTCTCGGGGTACAACAGGATGCCAGCTCAGCTGATATCCGCAAAGCC
TATCGGAAATTGTCCCTCACACTGCATCCCGATAAGAACAAAGACGAGAAT
GCCGAAACCCAGTTCAGACAGCTCGTGGCCATTTACGAGGTCCTGAAAGAC
GATGAGAGGAGACAGAGGTATGACGACATTCTGATCAACGGACTGCCCGAC
TGGCGACAGCCTGTGTTCTACTATAGGCGAGTGCGGAAGATGTCCAATGCC
GAGCTTGCCCTGCTTCTCTTCATCATTCTCACGGTTGGCCACTATGCAGTCGT
GTGGAGCATCTACCTGGAGAAACAGCTGGATGAGCTCCTGTCACGCAAGAA
GCGCGAAAAGAAGAAGAAAACCGGCAGTAAGAGCGTGGATGGGAAACCCA
TACCCAATCCACTCCTGGGTCTTGACTCTACTGGAGGGAGTGGCGGC

**Sec63(1-240)-V5**

GCCACCATGGCCGGACAACAGTTCCAGTATGACGACTCAGGGAACACCTTC
TTCTACTTCCTGACATCCTTCGTCGGCCTGATCGTCATACCAGCGACCTACT
ACCTCTGGCCCAGAGATCAGAACGCAGAACAGATACGGCTCAAGAACATTC
GTAAGGTGTATGGACGGTGCATGTGGTACCGCCTTAGACTGCTGAAACCTC
AACCCAACATCATACCCACTGTCAAGAAAATCGTGCTGTTGGCAGGATGGG
CTCTGTTTCTCTTTCTGGCCTACAAAGTGAGCAAGACAGACAGGGAGTATCA
GGAGTACAACCCGTATGAAGTGCTCAATCTTGATCCAGGCGCTACAGTGGC
CGAAATCAAGAAGCAGTATCGATTGTTGAGCCTCAAGTATCACCCCGATAA
AGGAGGCGATGAGGTGATGTTTATGCGCATTGCCAAAGCGTATGCAGCACT
GACTGACGAAGAGTCTAGGAAGAATTGGGAGGAGTTTGGGAATCCTGATGG
TCCACAAGCTACCTCCTTTGGTATCGCCCTTCCTGCTTGGATCGTTGACCAG
AAGAACAGTATTCTGGTCCTGCTGGTTTACGGACTTGCCTTTATGGTGATTC

TGCCAGTAGTAGTTGGGTCATGGTGGTATAGGTCCATCAGATACAGCGGGG
ACCAGATTCTGATTCGGACGACACAGATTTACACCTACTTCGTGTACAAAAC
CCGCAATGGCAAACCTATCCCGAATCCCCTGTTGGGGCTGGATAGTACTGG
CGGTTCTGGCGGC

**Tom20(1-33)-FLAG**

GCCACCATGGTGGGCAGGAACAGCGCCATCGCCGCCGGCGTGTGCGGCGCC
CTGTTCATCGGCTACTGTATCTATTTTGACCGGAAGCGGAGATCCGACCCCA
ATGATTACAAGGACCACGATGGCGACTATAAGGATCACGACATCGATTACA
AGGACGATGACGATAAGGGCGGCTCTGGCG

**Tom70(1-70)-3XFLAG**

GCCACCATGGCCGCGTCCAAACCCGTGGAAGCAGCCGTCGTTGCAGCCGCT
GTGCCGAGCTCTGGGTCAGGCGTGGGTGGAGGAGGAACCGCTGGTCCCGGA
ACAGGCGGCCTTCCTCGGTGGCAGTTGGCCCTGGCTGTAGGGGCACCACTG
CTCCTGGGAGCTGGGGCCATCTATCTGTGGAGCCGCCAACAGCGTCGAAGG
GAGGCCAGAGGGGACTATAAGGACCATGATGGGGATTACAAGGATCACGA
CATTGACTACAAGGATGACGACGATAAAGGCGGTAGTGGCGGC

**Table S1. Plasmids used in this study.**

| Plasmid code | Plasmid name | Promoter/terminator | Gene cloned | Referece |
|---|---|---|---|---|
| pYC1 | pRS316-Mmm1 | Own/Own | Mmm1 | [34] |
| pYC1 | pRS316-Mdm12 | Own/Own | Mdm12 | [34] |
| pYC91 | pRS314-Vps13-D716H | Own/Own | Vps13-D716H | [7] |
| pYM21 | pRS424-Mcp1 | Own/Own | Mcp1 | [14] |
| pYU41 | pRS313-GPDp-MCS-CYC1ter | GPD/CYC1 | None | This study |
| pYU47 | pRS314-GPDp-MCS-CYC1ter | GPD/CYC1 | None | This study |
| pYU53 | pRS315-GPDp-MCS-CYC1ter | GPD/CYC1 | None | This study |
| pYU54 | pRS315-ADH1p-MCS-CYC1ter | ADH1/CYC1 | None | This study |
| pYU59 | pRS315-ADH1p-MCS-CYC1ter | GPD/CYC1 | None | This study |
| pYU101 | pFA6a-mScarlet-KanMX4 | None/ADH1 | mScarlet | This study |
| pSFL1 | pTAC2-GFP1-10 | None | GFP1-10 | This study |
| pSFL2 | pTAC2-GFP11 | None | GFP11 | This study |
| pSFL9 | pRS316-GPDp-GFP1-10 | GPD/CYC1 | GFP1-10 | This study |
| pSFL10 | pRS314-GPDp-GFP1-10 | GPD/CYC1 | GFP1-10 | This study |
| pSFL11 | pRS316-GPDp-GFP11 | GPD/CYC1 | GFP11 | This study |
| pSFL12 | pRS314-GPDp-GFP11 | GPD/CYC1 | GFP11 | This study |
| pSFL15 | pRS314-GPDp-YNR021W-GFP11 | GPD/CYC1 | YNR021W-GFP11 | This study |
| pSFL16 | pRS316-GPDp-Ifa38-GFP1-10 | GPD/CYC1 | Ifa38-GFP1-10 | This study |
| pSFL17 | pRS315-GPDp-Ifa38-GFP1-10 | GPD/CYC1 | Ifa38-GFP1-10 | This study |
| pSFL19 | pRS316-GPDp-Vph1-GFP1-10 | GPD/CYC1 | Vph1-GFP1-10 | This study |
| pSFL20 | pRS315-GPDp-Vph1-GFP1-10 | GPD/CYC1 | Vph1-GFP1-10 | This study |
| pSFL21 | pRS314-GPDp-Vph1-GFP11 | GPD/CYC1 | Vph1-GFP11 | This study |
| pSFL22 | pRS316-GPDp-Tom71-GFP1-10 | GPD/CYC1 | Tom71-GFP1-10 | This study |
| pSFL23 | pRS314-GPDp-Tom71-GFP11 | GPD/CYC1 | Tom71-GFP11 | This study |
| pSFL26 | pRS316-GPDp-Pex3N-GFP1-10 | GPD/CYC1 | Pex3N-GFP1-10 | This study |
| pSFL28 | pRS316-GPDp-Erg6-GFP1-10 | GPD/CYC1 | Erg6-GFP1-10 | This study |
| pSFL61 | pRS316-GPDp-Ifa38-mCherry-GFP11 | GPD/CYC1 | IFA38-mCherry-GFP11 | This study |
| pSFL66 | pRS314-GPDp-Ifa38-3xFLAG-GFP11 | GPD/CYC1 | Ifa38-3xFLAG-GFP11 | This study |

| | | | | |
|---|---|---|---|---|
| pSFL67 | pRS314-GPDp-Erg6-3xFLAG-GFP11 | GPD/CYC1 | Erg6-3xFLAG-GFP11 | This study |
| pSFL69 | pRS314-GPDp-Pex3N-3xFLAG-GFP11 | GPD/CYC1 | Pex3N-3xFLAG-GFP11 | This study |
| pSFL73 | pRS314-GPDp-3xFLAG-GFP11 | GPD/CYC1 | 3xFLAG-GFP11 | This study |
| pSFL74 | pRS314-GPDp-V5-GFP11 | GPD/CYC1 | V5-GFP11 | This study |
| pSFL75 | pRS314-GPDp-Tom71-V5-GFP11 | GPD/CYC1 | Tom71-V5-GFP11 | This study |
| pSFL84 | pRS313-GPDp-Tom71-V5-GFP11 | GPD/CYC1 | Tom71-V5-GFP11 | This study |
| pFL16 | pRS315-GPDpBipN-mCherry-HDEL | GPD/CYC1 | BipN-mCherry-HDEL | This study |
| pFL24 | pRS315-ADH1p-mCherry-PTS1 | ADH1/CYC1 | mCherry-PTS1 | This study |
| pFL72 | pRS315-ADH1p-Erg6-mCherry | ADH1/CYC1 | Erg6-mCherry | This study |
| pMM73 | pCDNA3.1-Sec63(1-240)-V5-eGFP | CMV/bGH poly(A) signal | Sec63(1-240)-V5-eGFP | This study |
| pMM75 | pCDNA3.1-Tom20(1-33)-FLAG-eGFP | CMV/bGH poly(A) signal | Tom20(1-33)-FLAG-eGFP | This study |
| pMM76 | pCDNA3.1-Tom70(1-70)-FLAG-eGFP | CMV/bGH poly(A) signal | Tom70(1-70)-FLAG-eGFP | This study |
| pMM77 | pCDNA3.1-ERdj1(1-200-)V5-eGFP | CMV/bGH poly(A) signal | ERj1(1-200-)V5-eGFP | This study |
| pMM80 | pCDNA3.1_Tom20_GFP(1-10) | CMV/bGH poly(A) signal | Tom20(1-33)-FLAG-GFP1-10 | This study |
| pMM82 | pCDNA3.1_Tom70_GFP(1-10) | CMV/bGH poly(A) signal | Tom70(1-70)-FLAG-GFP1-10 | This study |
| pMM87 | pCDNA3.1_ERj1_GFP(11) | CMV/bGH poly(A) signal | ERj1(1-200-)V5-GFP11 | This study |
| pMM89 | pCDNA3.1_Sec63_GFP(11) | CMV/bGH poly(A) signal | Sec63(1-240)-V5-GFP11 | This study |

### Table. S2. Primers used in this study.

| Name | sequence (5'-3') |
|---|---|
| #YU291 | AATTGCGGCCGCATGTCTAGTTCAATATTTGGCCC |
| #YU292 | CCCGGATCCCTGAAATCTTGTTCTTTGCTTGTTTTTG |
| #YU293 | AATTGCGGCCGCATGACTTTTATGCAACAGCTTC |

| | |
|---|---|
| #YU294 | CCCGGATCCTTCCTTTTTAACCTGTCTTG |
| #YU295 | AATTGCGGCCGCATGGCAGAGAAGGAGGAAGC |
| #YU296 | CCCGGATCCGCTTGAAGCGGAAGAGCTTGC |
| #YU297 | AATTGCGGCCGCATGGCCGAAAACTCCCTCCTG |
| #YU298 | CCCGGATCCAAGCATGCCTTTAGCCCTATAAC |
| #YU305 | AATTGCGGCCGCATGGCCCCAAATCAAAGATC |
| #YU306 | CCCGGATCCGATCTGTTCTTTGATGAAGTG |
| #YU307 | AATTGCGGCCGCATGAGTGAAACAGAATTGAG |
| #YU308 | CCCGGATCCTTGAGTTGCTTCTTGGGAAG |
| #NU892 | AAAGCGGCCGCATGTTTTTCAACAGACTAAG |
| #NU893 | CCCACTAGTTTCTACATCATCGGCACCTCT |
| #NU946 | CCCACTAGTATGGTGAGCAAGGGCGAGGAG |
| #NU948 | AAACTCGAGTTACAATTCATCATGCTTGTACAGCTCGTCCATGCC |
| #YU377 | CCCGGATCCATGGTGAGCAAGGGCGAGGAGGAT |
| #YU378 | CCCGAATTCTTACAATTTTGAGCCACCAGACCCTCCCTTGTACAGCTCGTCCATGCC |
| #NU539 | CCCACTAGTATGGTGAGCAAGGGCGAGGAGGATAAC |
| #NU540 | CCCCGGATCCTTGTACAGCTCGTCCATGCCGCCGG |
| #YU1006 | CATAGGATCCTTACTTTTCGTTGGGATCTTTCGAA |
| #YU1007 | CATAGGATCCTTATGTAATCCCAGCAGCATTTACG |
| #YU1008 | CCCTCTAGAGCCACCATGGGTGGCACTAG |
| #YU1009 | CCCTCTAGAGCCACCATGGATGGAGGGTC |
